# Visualizing and quantifying data from timelapse imaging experiments

**DOI:** 10.1101/2021.02.24.432684

**Authors:** Eike K. Mahlandt, Joachim Goedhart

## Abstract

One obvious feature of life is that it is highly dynamic. The dynamics can be captured by movies that are made by acquiring images at regular time intervals, a method that is also known as timelapse imaging. Looking at movies is a great way to learn more about the dynamics in cells, tissue and organisms. However, science is different from Netflix, in that it aims for a quantitative understanding of the dynamics. The quantification is important for the comparison of dynamics and to study effects of perturbations. Here, we provide detailed processing and analysis methods that we commonly use to analyze and visualize our timelapse imaging data. All methods use freely available open-source software and use example data that is available from an online data repository. The step-by-step guides together with example data allow for fully reproducible workflows that can be modified and adjusted to visualize and quantify other data from timelapse imaging experiments.

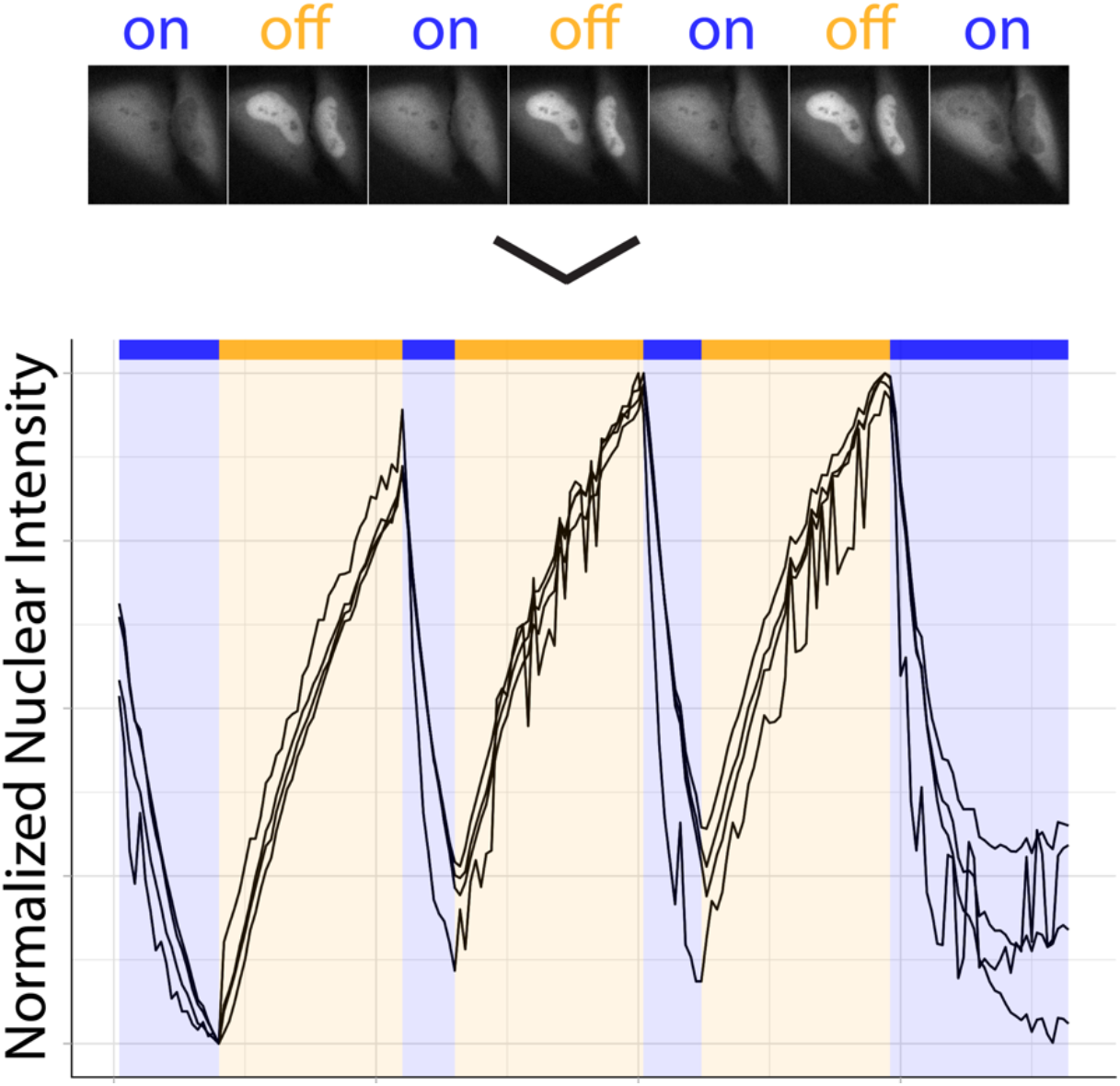

## 1 Introduction

An image is a representation of numbers in two dimensions. The simplest image that is acquired by fluorescence microscopy shows the distribution of fluorescence intensities in two spatial dimensions (x and y). However, it is very common that the output from an imaging setup is more complicated than a single image. This adds more dimensions to the imaging data. The two best-known examples of experimental data that has more than two dimensions is data from timelapse imaging, adding the dimension time and data from 3D-imaging, adding another spatial dimension. The addition of spectroscopic information (wavelength, fluorescence lifetime, anisotropy, photobleaching rate) brings yet another dimension to the imaging data. In box 1, several types of multidimensional datasets that are commonly encountered in fluorescence imaging are listed. Anything beyond a 2-dimensional (2D) dataset poses a data visualization challenge. Here, we will specifically address data from timelapse imaging. Representing data with 3 spatial dimensions (3D imaging) as obtained from confocal imaging, 3D-STORM or light-sheet microscopy needs a different approach [1,2] and will not be discussed here.

#### Box 1

Examples of common types of multidimensional data acquired by fluorescence imaging. Here, x,y,z are spatial dimensions, t is time, λ a spectral dimension and τ is lifetime.

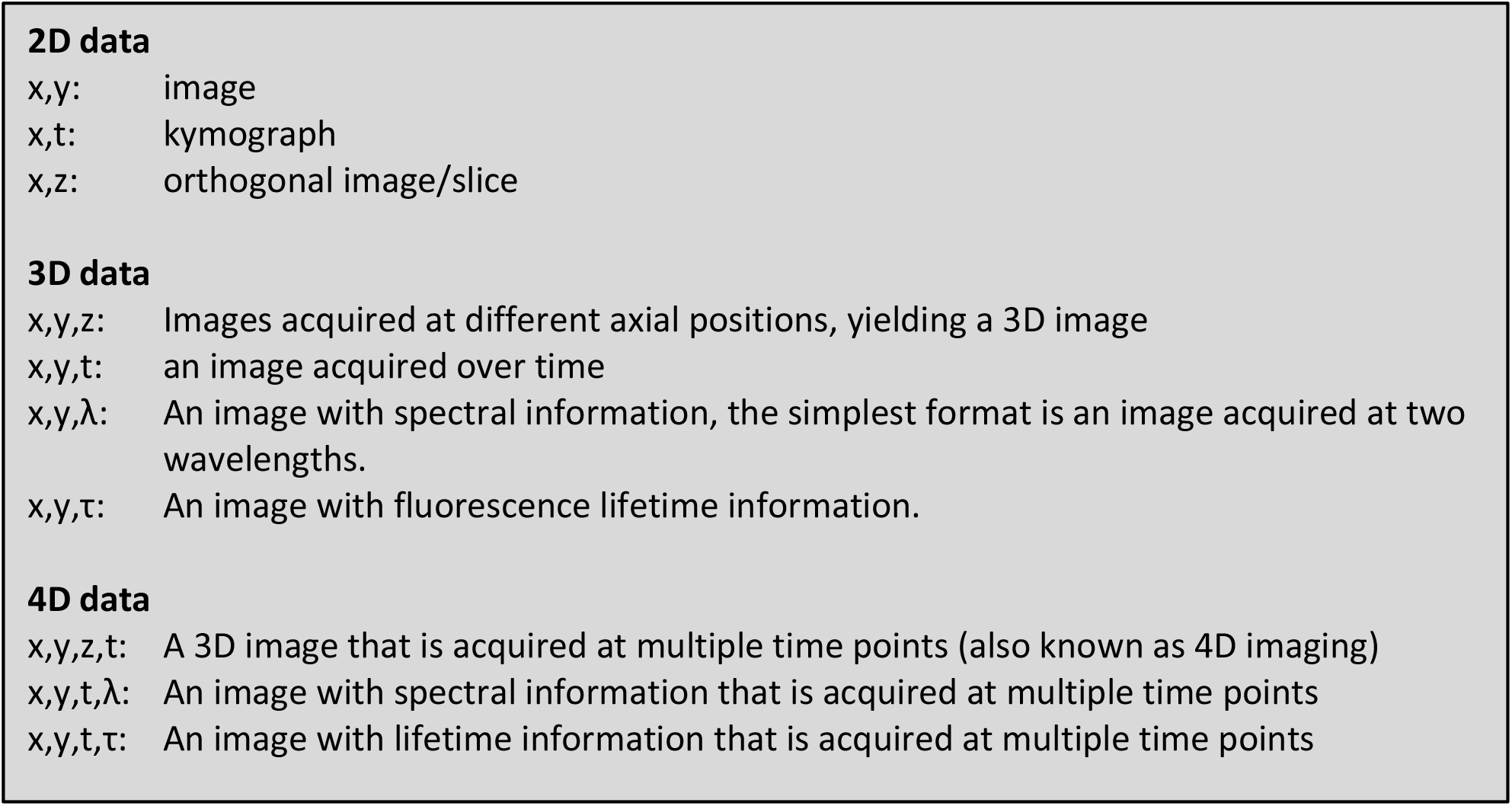

The choice for the type of data visualization depends on the information that is extracted from the data. Below, we explain some of the data visualizations that we use in our research. This document is formatted as a ‘protocol’ with the necessary materials listed in section 2, the protocols in section 3 and notes in section 4.

The materials in section 2 comprise (open source) software, code and data and are all freely available online. The step-by-step guides in section 3 should be self-explanatory and are tested on MacOS and Windows operating systems. Since it is impossible to test the software and protocol for each possible computer configuration, you may need to update software or tweak the protocol to make it work on your system. If you run into problems and find a solution, we would be happy to hear about it as it will help us to improve the workflows. The expected outcome for each of the workflows is shown as a figure or plot at the end of the protocol. In general, the interpretation and biological significance of the result can be found in the papers that we refer to in the introduction of each workflow.

## 2 Materials

### 2.1 Mapping dynamics of structures

- FIJI; https://fiji.sc [3]. We use text between brackets to indicate the sequence of menu items that needs to be selected to reach a function. For instance [File > New > Image…] to create a new image (Note 1).
- LUTs that are used here are available in the Github repository: https://github.com/JoachimGoedhart/TimeLapse-Imaging-DataViz
- To install a LUT, open the folder with LUTs: [File > Show Folder > LUTs]. Copy the .lut files to this folder. The LUTs are available after restarting FIJI.
- Example data; https://doi.org/10.5281/zenodo.4501412

### 2.2 Plotting signals from timelapse imaging

- FIJI; https://fiji.sc [3]
- Set the measurements to only measure mean gray values: [Analyze > Set Measurements…] and select ‘Mean gray value’, ‘Display label’ and set ‘Decimal places’ to 2.
- Macros that are used here are available in the Github repository: https://github.com/JoachimGoedhart/TimeLapse-Imaging-DataViz
- To install a macro use [Plugins > Macro > Install…] and locate the .ijm file. The macro will appear in the menu [Plugins > Macro]. After closing FIJI, the macro is removed from the menu. If you want to use the macro more often, you can add it to the plugins menu: [File > Show Folder > Plugins]. Move the .ijm file into this folder. After restarting FIJI, the macro is available from the Plugins menu.
- Example data; https://doi.org/10.5281/zenodo.4501412
- PlotTwist; https://huygens.science.uva.nl/PlotTwist [4]
- R; https://www.r-project.org/
- R package ‘tidyverse’ (https://tidyverse.tidyverse.org). The package can be installed in Rstudio from the menu: [Tools > Install Packages…]
- Rstudio; https://rstudio.com/products/rstudio

## 3 Methods

### 3.1 Mapping dynamics of structures

The acquisition of fluorescence images at defined time points can be used to examine the dynamics of objects in the images. There are several ways to do this and four relatively straightforward methods are described below. The methods described here are used to visualize the dynamics of cells and within cells. More advanced methods can be found elsewhere, for instance tracking of objects [5] or by measuring correlations between subsequent frames [6].

#### 3.1.1 Encoding dynamics with temporal color coding

A very straightforward way of visualizing the dynamics in an image format is by applying unique colors to each frame of a movie. In this way, the color encodes time. This method works very well for showing the trajectory of objects in an image. In this example, we use temporal color coding to display the migration of white blood cells that are labeled with a fluorophore on a 2D monolayer of endothelial cells.

- Open the file ‘201029_PMN_channel_06-2.tif’ in Fiji by dropping it onto the main window (Note 2).
- [Image > Hyperstacks > Temporal-Color Code] (Note 3)
- Choose a LUT, here we selected ‘Fire’. Make sure to select the box ‘Create Time Color Scale Bar’. (Note 4)
- Two images will be displayed, one with the color coded timelapse and another with the LUT, showing which frames are represented by the different colors.

For the final result, we copy the LUT and paste it in the image ‘MAX_colored’:

- Activate the ‘color time scale’ window and select the contents: [Edit > Selection > Select All].
- Copy the selected contents, [Edit > Copy], and paste in the image ‘MAX_colored’: [Edit > Paste]. Move the LUT to the right location. Deselecting it (by clicking next to the ROI) fixes the location.

**Figure 1:**
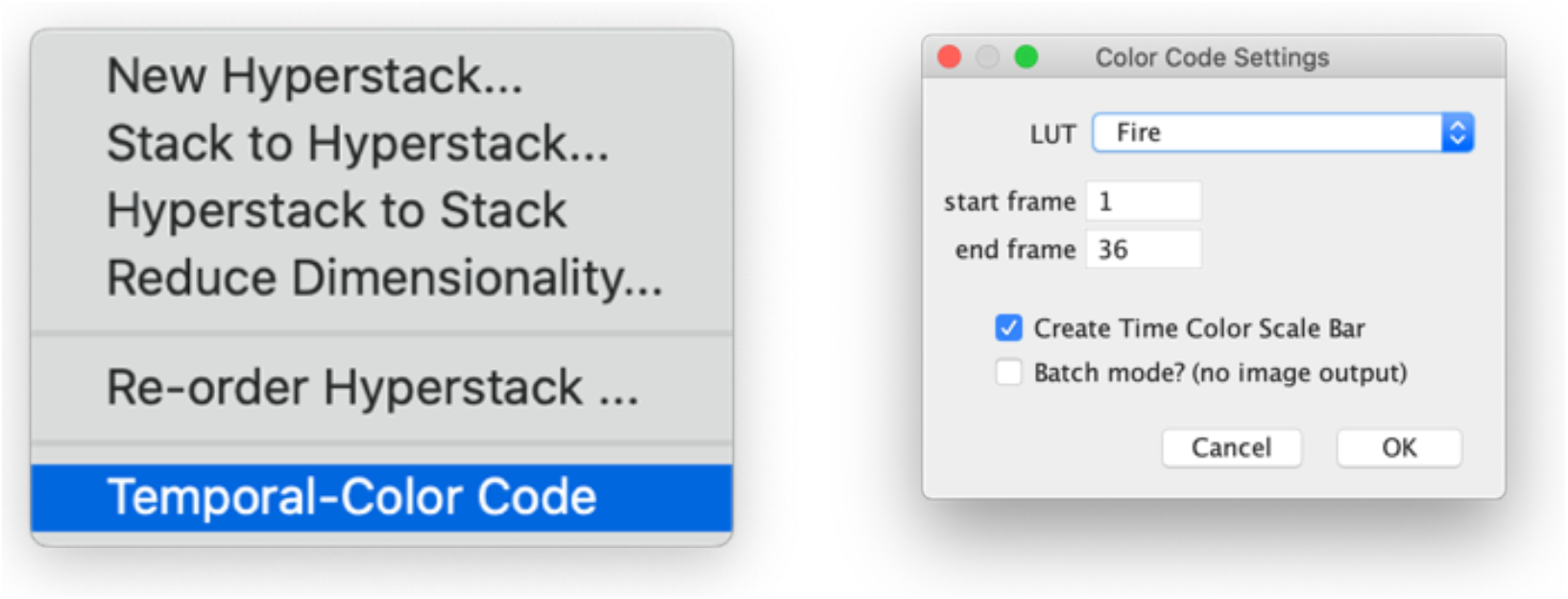
Screenshots of a submenu and the window that pops up when the ‘Temporal-Color Code’ is selected.

**Figure 2:**
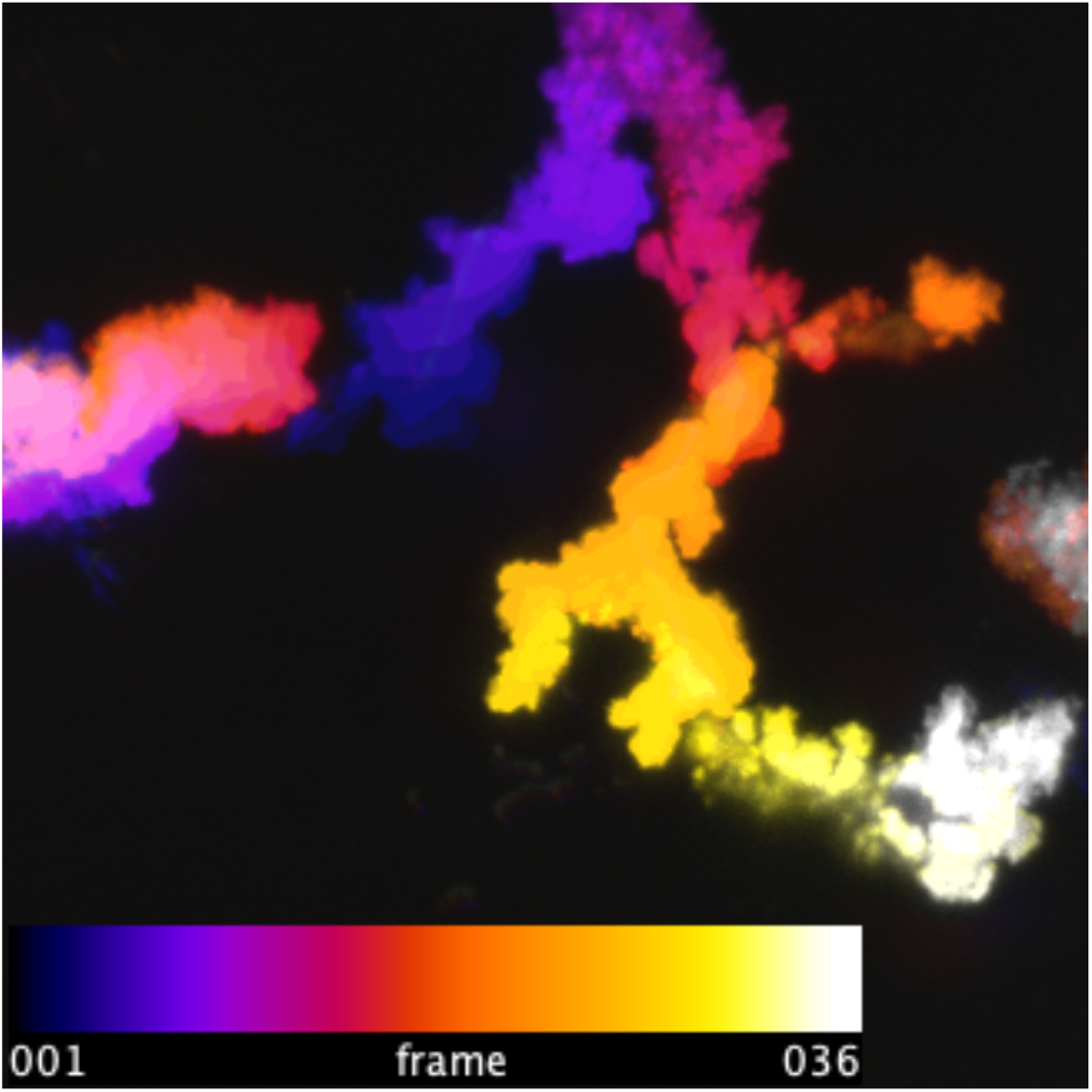
Migration of a white blood cell visualized by applying the temporal color code. In this figure the ‘Fire’ look-up table was used to define the colors for the different frames.

#### 3.1.2 Kymographs

In image analysis, a kymograph is an image in which one dimension is space and the other dimension is time. Kymographs are used to visualize the movements of structures or organelles. An image with one spatial and one temporal dimension can be directly generated by timelapse imaging of a line (instead of a frame) with a confocal microscopy. But usually, kymographs are created from movies that are obtained by timelapse imaging. To this end, a line is defined in the image by the user, which is used to create the kymograph. Here, we use a kymograph to analyze the dynamics of microtubules that are labeled at their tips with fluorescently tagged EB3 [7]. Midway through the timelapse, nocodazole is added to disrupt microtubule integrity and dynamics.

- Locate the image sequence ‘EB3-timelapse.tif’ on your computer and open it in FIJI.
- Alternatively, this file can be retrieved through a URL [File > Import > URL…] with https://github.com/JoachimGoedhart/TimeLapse-Imaging-DataViz/raw/main/3_1_2_KymoGraph/EB3-timelapse.tif
- Draw a line in the image, using the line tool from the main window in FIJI (Note 5)

**Figure 3:**
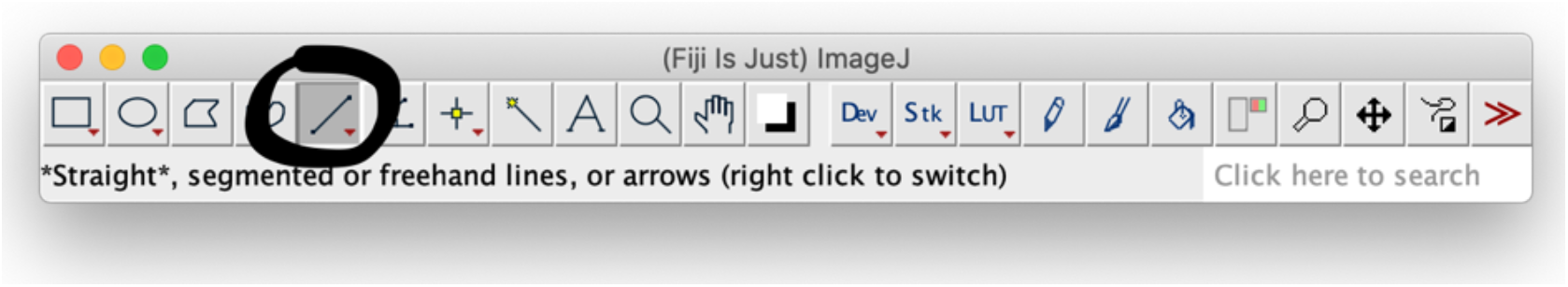
Screenshots of the main window of FIJI with the line tool selected and highlighted by a black circle.

- Retrieval of the ROI that was used here can be done by drag and drop of the file onto the main window of Fiji.
- To reproduce the line that was used in this example, drop the file ‘Line-for-Kymograph.roi’ onto the main window.
- Generate a kymograph: [Analyze > Multi Kymograph > Multi Kymograph], Linewidth 1
- Stretch the image in the time dimension to improve the visualization: [Image > Scale…] with ‘Y Scale:’ set to 4 (and ‘X Scale:’ to 1). (Note 6)

**Figure 4:**
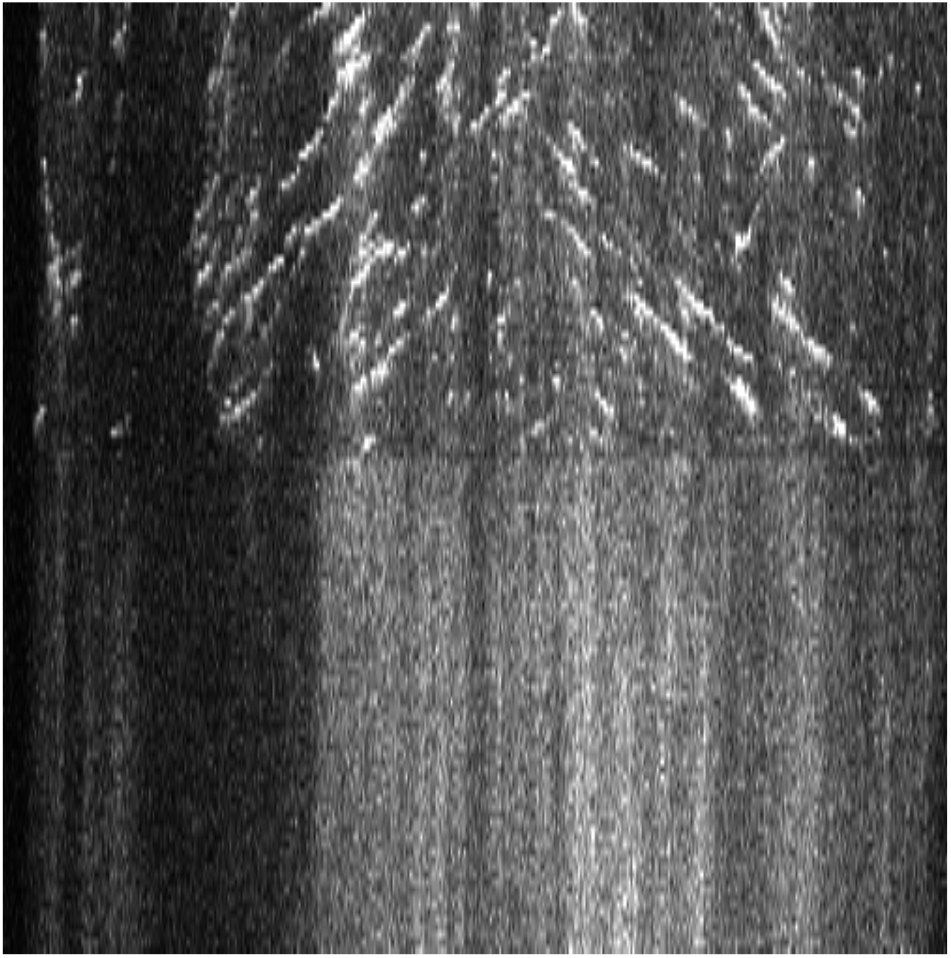
A kymograph of microtubule dynamics, in which moving tips are visible as diagonal lines. Time progresses in the vertical direction from the top to the bottom and the horizontal direction is a spatial dimension (reflecting the selected line).

- To save the original image with the colored line that was selected to generate the kymograph, first convert the data to allow the display of color: [Image > Type > RGB color]
- Add the selected line to the ROI manager: [Analyze > Tools > ROI Manager…] and press the button ‘Add’.
- To make the line more visible, select the ROI from the list and press ‘Properties’ and change the Width: 2.
- To show the line in the image use the button [More » Fill]
- To use only this frame from the stack use [Image > Duplicate…] and deselect ‘Duplicate stack’. (Note 7)
- Save the image as a .PNG file.

**Figure 5:**
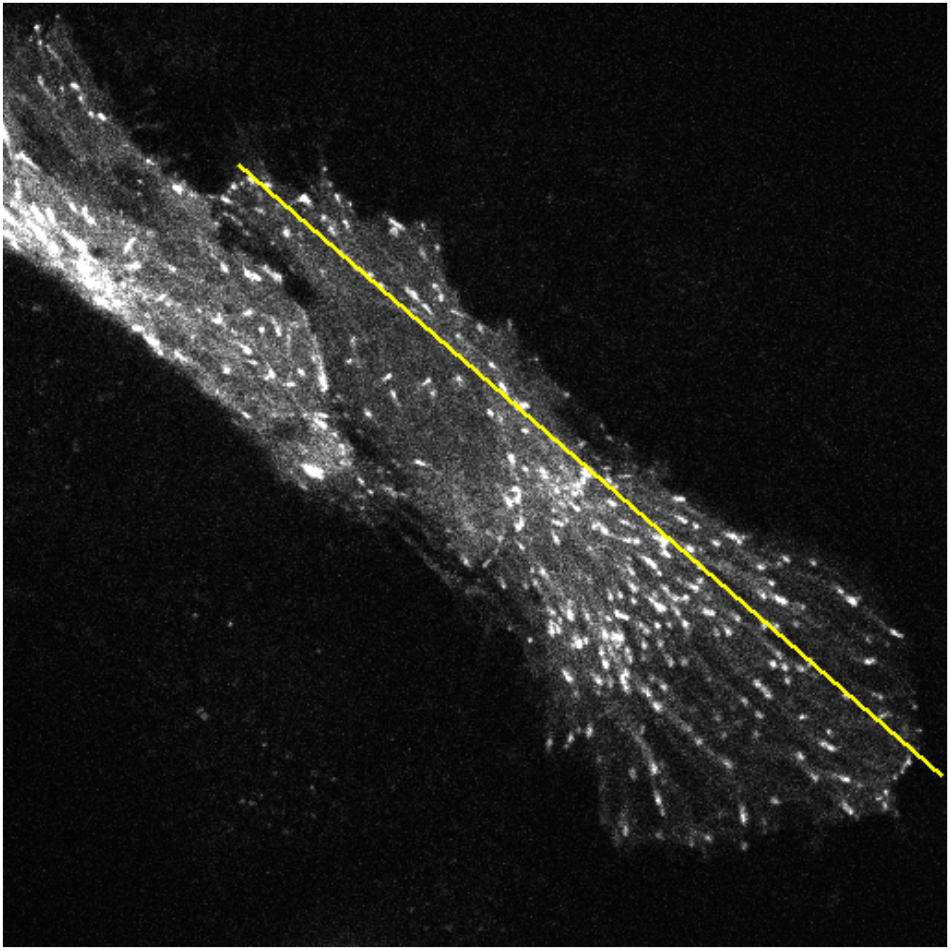
A still image from a timelapse imaging dataset that shows the line (in yellow) from which the kymograph (shown in figure 4) was generated.

#### 3.1.3 Visualizing fluctuations in intensity

We have recently explored another way to visualize dynamics in data acquired by timelapse imaging [8]. In this case, we wanted to quantify how intensities of fluorescence signals changes over time by determining the coefficient of variation (standard deviation divided by the mean) on a pixel-by-pixel basis. The imaging data was acquired from endothelial cells of which the plasma membrane was labeled with the fluorescent protein mNeongreen tagged with a CaaX box [8].

- Open the sequence ‘mNeon-CAAX.tif’
- Any other sources of fluctuations or changes in intensity are corrected for. Image drift can be corrected as described for ‘<u>Visualizing a change in area</u>’ and is not necessary here. Intensity loss due to bleaching is corrected by [Image > Adjust > Bleach Correction] with ‘Correction Method:’ set to ‘Exponential Fit’
- The resulting image ‘DUP_mNeon-CAAX.tif’ is used to determine the standard deviation in groups of 10 images. For instance, if we have 100 images, these are grouped in batches of 1-10, 11-20, 21-30, etc. In FIJI this is achieved as follows: [Image > Stacks > Tools > Grouped Z Project…]
- Choose ‘Standard Deviation’ as the projection method and group size 10.

**Figure 6:**
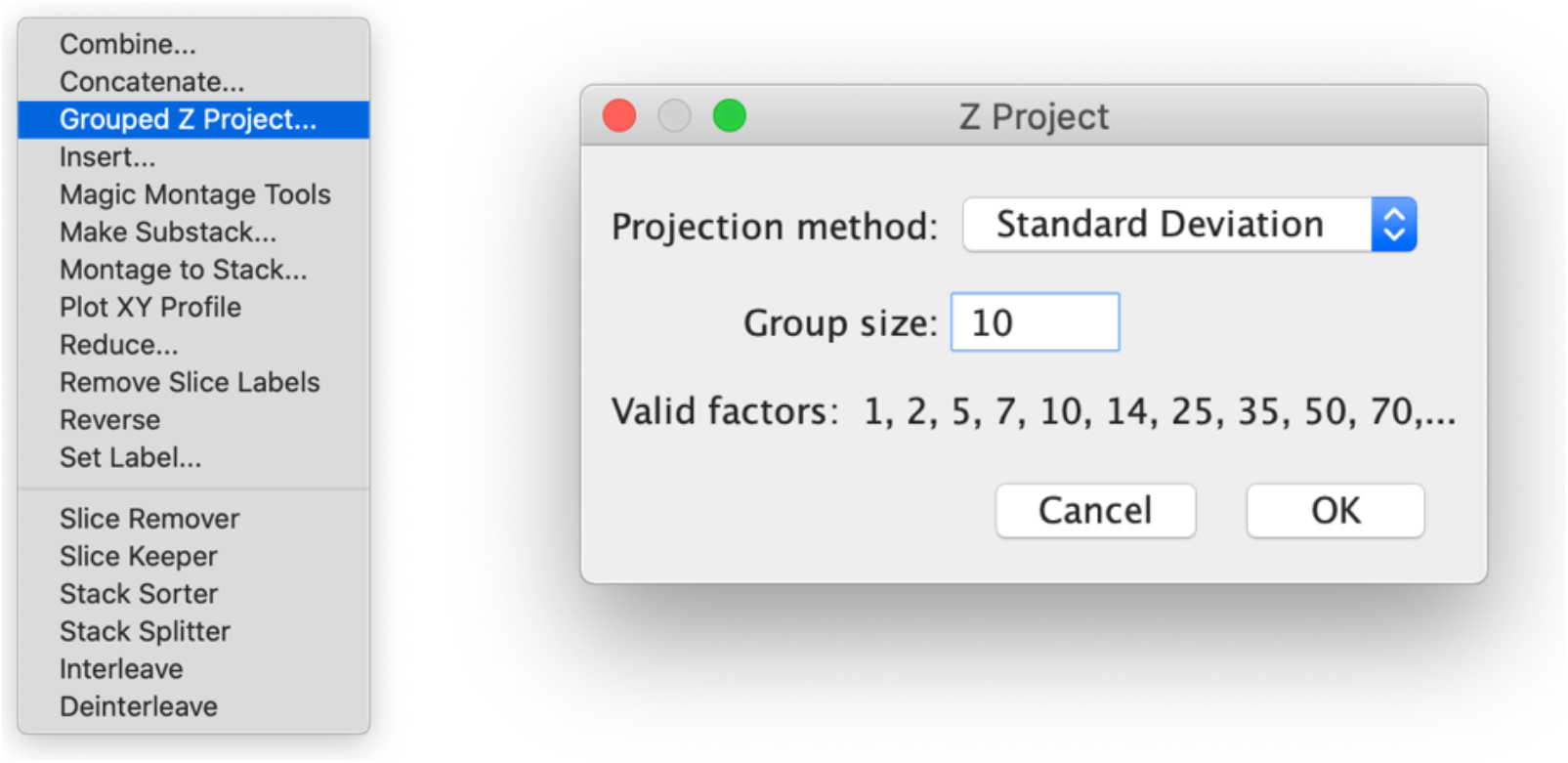
Screenshots of a submenu and the window that pops up when the ‘Gropuped Z Project…’ is selected.

- On the same sequence, the average per 10 frames is calculated: select the window ‘DUP_mNeon-CAAX.tif’. Then select [Image > Stacks > Tools > Grouped Z Project…] and choose ‘Average Intensity’ as the projection method and group size 10.
- Divide the stack with standard deviation data by the stack with average intensities by using [Process > Image Calculator…] and selecting Image1: STD_DUP_mNeon-CAAX.tif, Operation: Divide and Image2: AVG_DUP_mNeon-CAAX.tif. Make sure that that both checkboxes are selected (‘Create new window’ and ‘32-bit (float) result’).

**Figure 7:**
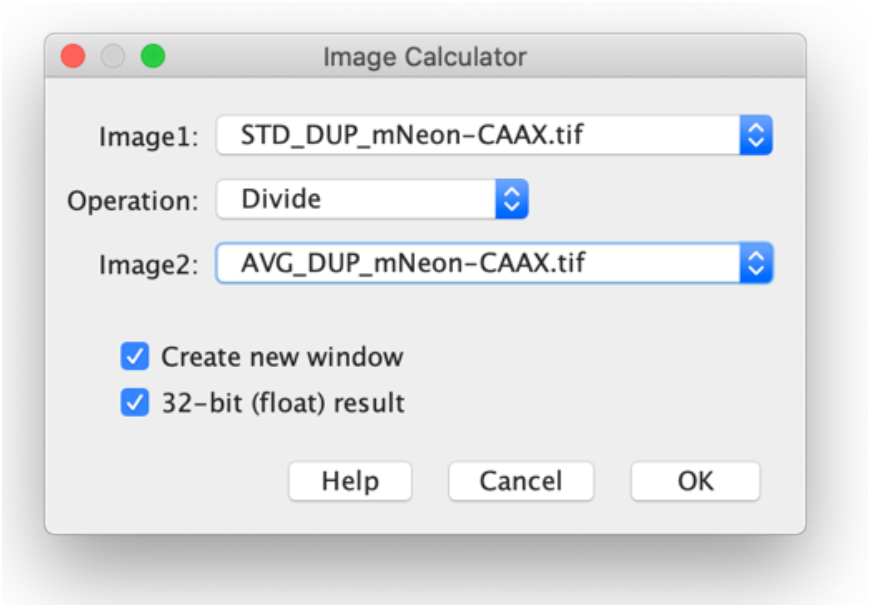
Screenshot of the image Calculator window with the settings that are used to divide the stack with the standard deviation of intensities over the stack with the average intensity.

- The resulting stack ‘Result of STD_DUP_mNeon-CAAX.tif’ shows how the intensity-averaged standard deviation changes over time. To generate an average image, choose [Image > Stacks > Z Project…] and ‘Project type’ is ‘Average Intensity’ (‘Start slice’ is 1 and ‘Stop slice’ is 35).
- a LUT can be applied to the map to better show the differences in dynamics: [Image > Lookup Tables]. Here, we have selected the LUT ‘morgenstemning’ [9] (See 2.1 on how to install this LUT).
- The image can be optimized by adjusting the brightness and contrast manually: [Image > Adjust > Brightness/Contrast…]
- To reproduce the image displayed here, use [Process > Enhance Contrast] with ‘Saturated pixels:’ set to 0.3 %.
- For compatibility with downstream applications (text editors, presentation software) it is useful to save the result in PNG format: [File > Save As > PNG…]

**Figure 8:**
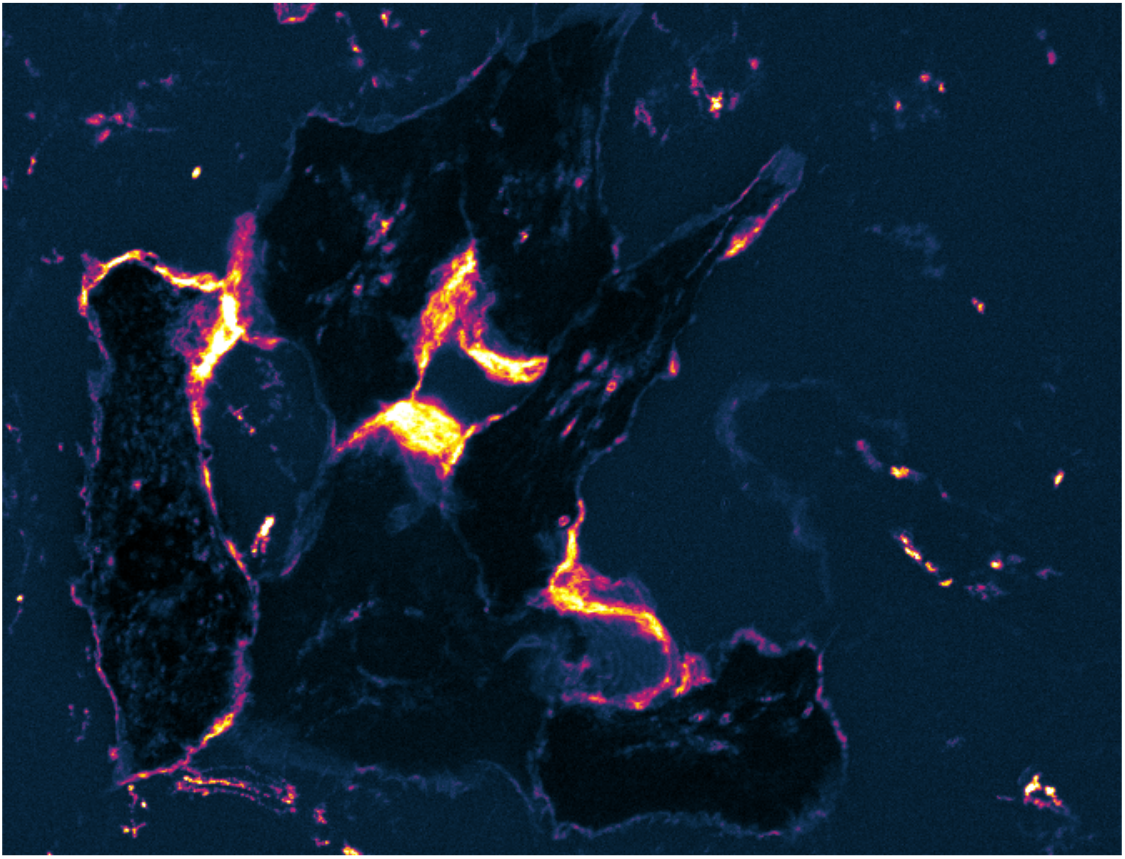
The result of the workflow that shows intensity fluctuations in a timelapse imaging dataset. The lighter the color, the larger the change in intensity during the timelapse. The light colors are interpreted as areas where the membranes are highly dynamic.

#### 3.1.4 Visualizing a change in area

Cells may change their shape in response to perturbations. Endothelial cells respond to thrombin with a Rho GTPase mediated cell contraction, which strongly reduces their cell area. To visualize the change in area between two (user selected) time points, we use a colormap that visualizes reduced, increased and unchanged cell area [10]. The data used is here is from an endothelial cell that expresses a membrane marker (mTurquoise2-CaaX). The change in cell area is triggered by treating the cells with thrombin [10,11].

- Open the image ‘2010230_BOEC-mTq2-CaaX.tif’ from your computer or use the URL: https://zenodo.org/record/4501412/files/2010230_BOEC-mTq2-CaaX.tif
- Remove image shift by using the plugin ‘Linear Stack Alignment with SIFT’: [Plugins > Registration > Linear Stack Alignment with SIFT] and use default settings.
- Crop the stack to exclude the cells at the edges, by drawing an ROI with the Rectangle tool and remove everything outside of the ROI: [Image > Crop]
- Get frame 4: [Image > Duplicate…]. Set the ‘Title’ to Frame4 and uncheck ‘Duplicate stack’
- Repeat to get frame 25, name this image Frame25
- [Process > Filters > Gaussian Blur…] and set Sigma(Radius) to 2
- [Image > Adjust > Threshold…] and use the button ‘Set’ to set the ‘Low threshold level’ to 270 (and the ‘High threshold level’ to 65535). After pressing the button ‘OK’, press the button ‘Apply’ on the Threshold window (make sure that the ‘Dark background’ box is checked).
- Repeat the previous step for image Frame25 with the same settings for thresholds
- Both images are now ‘binary’ images with values of either 0 (background) or 255 (foreground). Use [Process > Binary > Fill Holes], to fill holes in both binary images.
- On image Frame4: [Process > Math > Subtract…], Value: 254
- On image Frame25: [Process > Math > Subtract…], Value: 253
- Sum the images: [Process > Image Calculator…], Image1: Frame4, Operation: Add, Image2: Frame25 and check ‘Create new window’
- The resulting image has pixel values of 0, 1, 2 and 3
- To apply a colormap to depict these different pixel values, take the file ‘AreaLUT.lut’ and drop this file on the main window. When this LUT is used, red depicts the area loss, blue is expanded area and white is unchanged between the two selected frames.
- The LUT can be modified to change the colors: [Image > Color > Edit LUT…]

**Figure 9:**
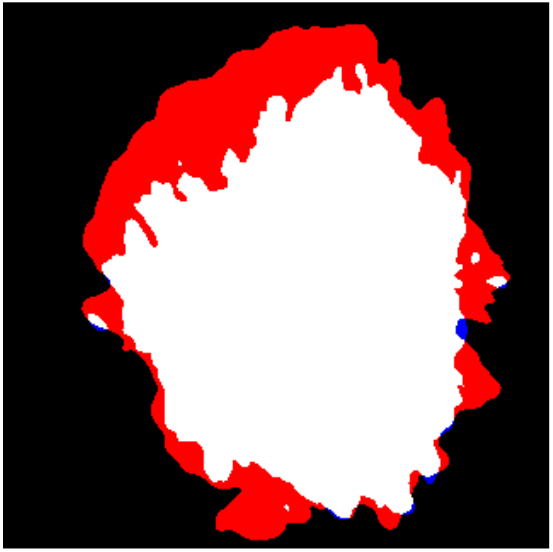
A four color image that visualizes a change in cellular area. The white area is unchanged, the red color indicates a loss of area and the blue color indicates a gain in area. As such, the red color visualizes cell retraction, and the blue color depicts cell expansion or protrusion.

### 3.2 Plotting signals from timelapse imaging

In our work, we use genetically encoded biosensors to report on intracellular processes. These biosensors may report on changes in calcium levels, kinase activity or lipid signaling. In addition, we used chemo- and optogenetic switches to steer cellular processes. In all of these cases, we analyze changes in optical signals over time. The signal can simply be the fluorescence intensity of a reporter, but it may also be the ratio of two intensities. Ratios of signals are more robust to photobleaching, cell shape changes and sample drift than an intensity measured from a single channel. In the next workflows, we will explain how intensities can be quantified and visualized over time.

#### 3.2.1 Visualizing the dynamics of intensity

The LEXY probe consists of a fluorescent protein and a light inducible nuclear export sequence [12]. In resting cells, the fluorescence is mainly located in the nucleus. When illuminated with blue light, the nuclear export signal is exposed, and the probe moves out of the nucleus. In absence of blue light, the system reverses to the original state and the probe accumulates in the nucleus. Here we quantify the nuclear intensity over time, to demonstrate the nuclear export induced by blue light.

- Open the data: ‘210203_LEXY_combined.tif’ from your computer or use the URL: https://zenodo.org/record/4501412/files/210203_LEXY_combined.tif
- Find an area with no fluorescence, which can be used as background. You may need to stretch the contrast to find the optimal background region: [Image > Adjust > Brightness/Contrast…]
- Draw an ROI that represents background signal
- Run the macro Subract_Measured_Background which will, for each image in the stack, subtract the average background value calculated from the ROI. See 2.2 for instructions on how to install the macro.
- Activate the ROI manager: [Analyze > Tools > ROI Manager…]
- Draw an ROI in the image sequence to select the nucleus of a cell, add to ROI manager ‘Add [t]’
- Repeat the selection of nuclei and the addition of ROIs to the ROI manager until all nuclei are marked
- Rename the ROIs: select a ROI from the list and use [Rename…] on the ROI manager. Use a more informative, here we use Cell-01, Cell-02, etc.
- Save the set of ROIs: [More » Save…]
- To reproduce the ROIs that are used in this example, drop the file ‘RoiSet.zip’ onto the main window.
- Select all ROIs in the ROI manager, except for the background ROI, if it is in the list (select the first label, then select the last label in the list, while pressing the <shift> key)
- Make sure that the measurements are correctly defined: [Analyze > Set Measurement] and select ‘Mean gray value’, ‘Display label’ and set ‘Decimal places’ to 2.
- Measure in all ROIs the mean gray value: [More » Multi Measure] and select ‘Measure all slices’ and make sure to that ‘One row per slice’ is not selected. (Note 8)
- Save the table with results in csv format: [File > Save As…].

- The open-source web application will be used to plot the data: https://huygens.science.uva.nl/PlotTwist/
- The PlotTwist app needs an internet connection. It disconnects when it is inactive for a while. Both issues can be addressed by running it offline from Rstudio, for instructions see: https://github.com/JoachimGoedhart/PlotTwist
- Select ‘Upload (multiple) file(s)’ and use ‘Browse…’ to locate the Results.csv file or drop the file on the ‘Browse…’ button to upload the data.
- The app splits the ‘Label’ column into three columns, of which the ‘Sample’ column is used to identify the different ROIs.
- Select the checkbox ‘These data are Tidy’.
- Set the x-axis variable to ‘Slice’ and the y-axis variable to ‘Mean’. The default settings for identifier of the sample and condition (‘Sample’ and ‘id’ respectively) can be used for these data.
- Optional: a data normalization can be applied in PlotTwist. Several options are available when the checkbox ‘Data normalization’ is activated. Here, we correct for differences in intensity due to different expression levels between cells. Select the option ‘Rescale between 0 and 1’.
- To visualize the data, click on the ‘Plot’ tab. A plot of the data is shown, and the plot can be modified to improve the visualization.
- To store the settings of the user interface, use the button ‘Clone current setting’ above the plot.
- The URL can be stored and used later to reproduce the exact settings. For instance, the URL that belongs to the plot shown here is: https://huygens.science.uva.nl:/PlotTwist/?data=3;TRUE;TRUE;zero_one;1,5;&vis=dataasline;1;;;1;;&layout=;;;;;;;6;X;480;600&color=none&label=TRUE;Lightinducednuclearexportinsinglecells;TRUE;Slice;Normalizednuclearintensity;TRUE;26;24;18;8;;;&stim=TRUE;both;1,20,20,55,55,65,65,101,101,112,112,148,148,182;on,off,on,off,on,off,on,off;blue,orange&
- To use this setting, copy-paste the URL in a browser or use this link. Upload the data and select the x-axis and y-axis variable. Click on the ‘plot’ tab and a plot that looks identical to the one shown here should appear.

**Figure 10:**
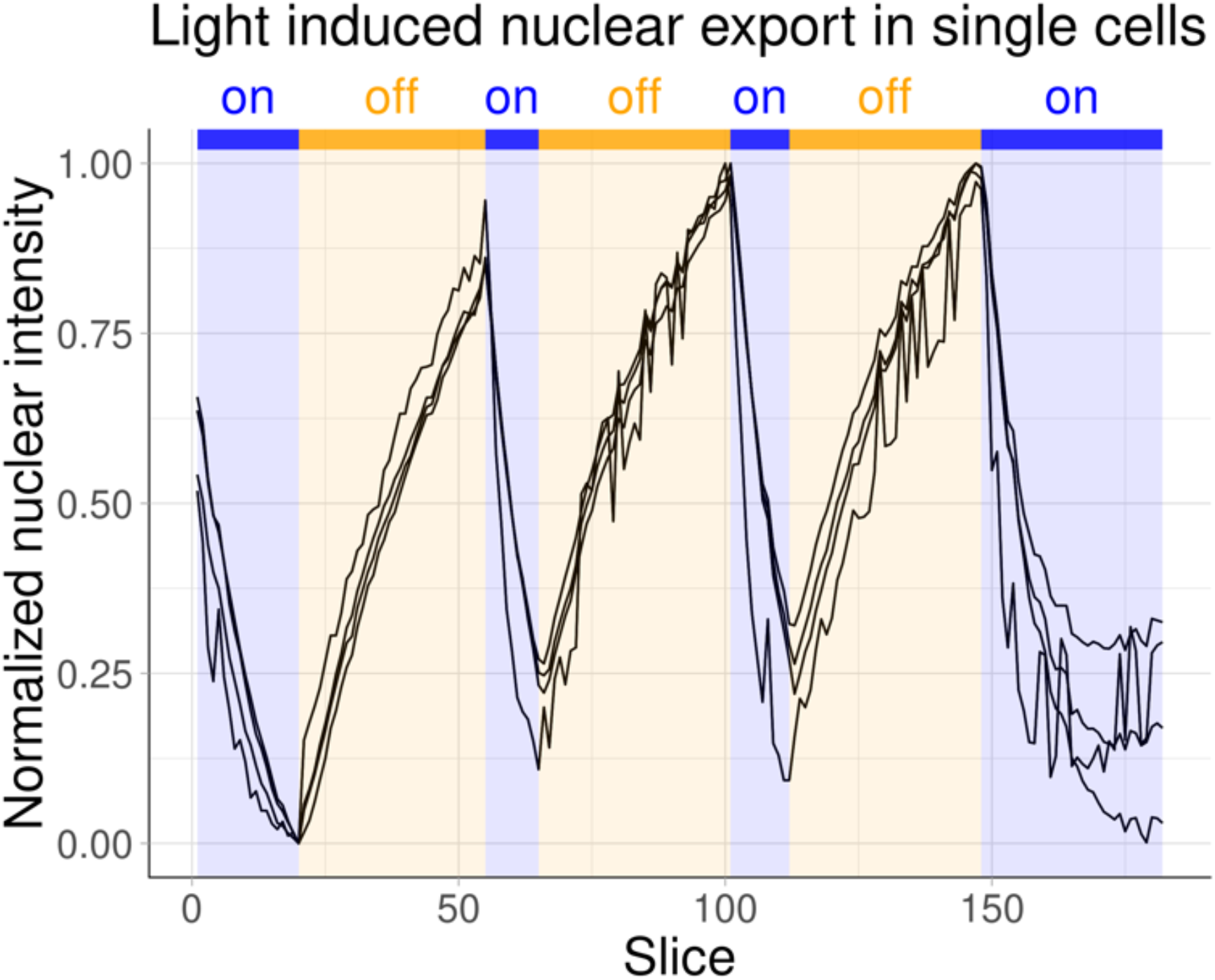
A plot that shows the change in nuclear intensity over time and how it depends on exposure to blue light. When the light is on, the nuclear intensity decreases and when the light is off, the intensity increases. The black lines represent the data from four different cells. The data was scaled between 0 and 1.

- A limitation of this workflow is that you end up with ‘Slices’ at the x-axis instead of time. This can be fixed with an R-script, as will be explained in section **3.2.2**. An alternative is to run an ImageJ macro ‘Add-timing-to-results.ijm’ (available in the repository on Github: https://github.com/JoachimGoedhart/TimeLapse-Imaging-DataViz/tree/main/Macros) that adds a column ‘Time’ to the results window that is calculated from the slice number and a user-defined time interval.

#### 3.2.2 Visualizing ratiometric data

Yellow cameleon is a FRET-based calcium biosensor [13]. The ratio of the intensity at two wavelengths is used as a read-out. When calcium levels are high, the cyan intensity is low and the yellow intensity is high and when calcium levels are low, this is reversed. Therefore, a plot of the ratio of yellow over cyan signal is often shown to visualize calcium dynamics over time. Here we used yellow cameleon to measure calcium oscillations triggered by histamine in HeLa cells. The histamine receptor is deactivated by the addition of the antagonist pyrilamine [14] near the end of the timelapse.

- The data consists of individual frames that are located in the folder YCam. The data is available here: https://zenodo.org/record/4501412/files/YCaM.zip
- [File > Import > Image Sequence…] and locate the folder ‘YCam’ with the images and press ‘open’. To open the data acquired from the CFP emission channel use for ‘File name contains:’ ‘CFP’ and make sure that the checkbox ‘Sort names numerically’ is selected.
- Rename the image sequence to CFP: [Image > Rename…]
- Repeat the previous two steps to get the YFP data.
- The background (or offset) is removed by determining the average background signal from a region in the image that does not contain signal from cells. First draw a ROI in a region that represent background. Run the macro ‘Subtract_Measured_Background’ (https://github.com/JoachimGoedhart/TimeLapse-Imaging-DataViz/tree/main/Macros)
- Repeat the background subtraction for the YFP image sequence.
- The images can be saved (and are available as CFP.tif and YFP.tif)
- ROIs that define individual cells can be drawn and add to the ROI manager (see also 3.2.1).
- To reproduce the ROIs, that are used in this example, drop the file ‘RoiSet.zip’ onto the main window.
Make sure that the measurements are correctly defined: [Analyze > Set Measurement] and select ‘Mean gray value’, ‘Display label’ and set ‘Decimal places’ to 2.
- To obtain the fluorescence intensity from multiple ROIs over time select the ROIs in the ROI manager. Next, apply [More » Multi Measure] and select ‘Measure all 121 slices’. Deselect the other two checkboxes. The resulting window with data can be saved: *File > Save As…* in csv format. The resulting file is named ‘Results-CFP.csv’
- Repeat this for the YFP data and name the file with results ‘Results-YFP.csv’.

At this point, the results can be inspected with the online visualization tool PlotTwist:

- Choose the option ‘Upload (multiple) file(s)’ and drag simultaneously the files Results-CFP.csv and Results-YFP.csv onto the ‘Browse…’ button to upload.
- Select the check box ‘These data are Tidy’
- Select the variables, for x-axis = Slice, y-axis = Mean, Identifier of samples = ‘Sample’ and Identifier of conditions =’id’.
- Activate the checkbox ‘Data normalization’ and use the default setting: ‘Fold change over baseline (I/I0)’
- Press on the ‘Plot’ tab to show the plot
- The default plot will show up and the visualization can be further optimized, if needed.

**Figure 11:**
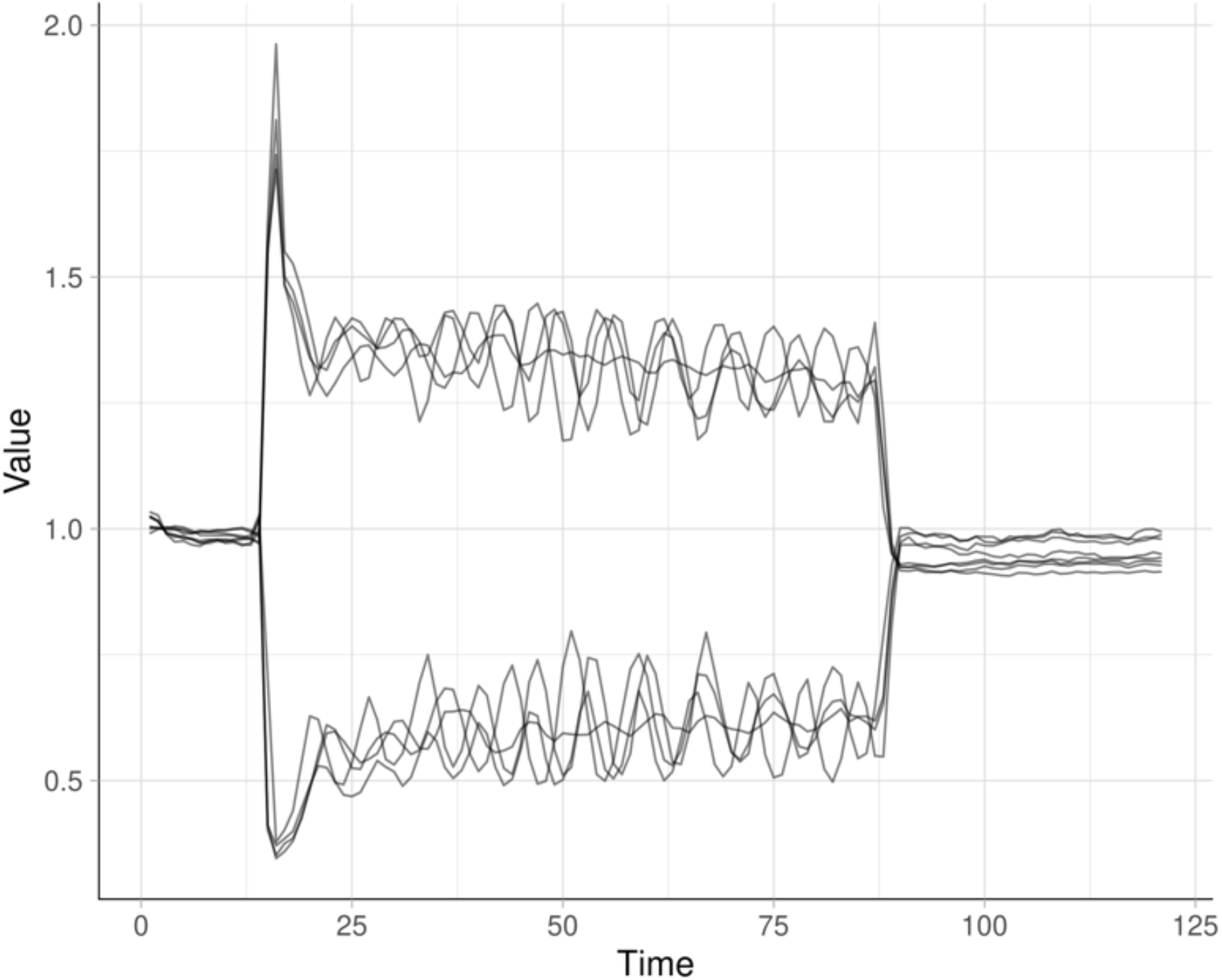
A plot that shows the change in fluorescence intensity over time from the CFP and YFP channel. The black lines represent the data from four different cells. The baseline fluorescence, determined from the average of the first 5 frames, was normalized 1.

Below, we explain how the data can be processed to normalize the data, calculate the ratios and visualize as a plot in R. See 2.2 for instructions on how to install R and R studio.

- Make sure that the R-script ‘Plot_Ratiometric-data’ and the csv files ‘Results-YFP’ and ‘Results-CFP’ are located in the same folder. Make sure to use these files names, otherwise the script does not find the data.
- Open the script in Rstudio by double clicking on the R-script.
- In Rstudio go to [Session > Set Working Directory > To Source File Location]
- The code can be run line-by-line to understand what each line of code does: Position in the line you want to run and press: <command>+<enter> or use the menu: [Code > Run Selected Line(s)]
- After running a line of code, the cursor moves down and so the previous step can be repeated until all the code is executed. (Note 9)
- The script will produce a plot, that is saved in the working directory as ‘Ratio-plot.png’.

**Figure 12:**
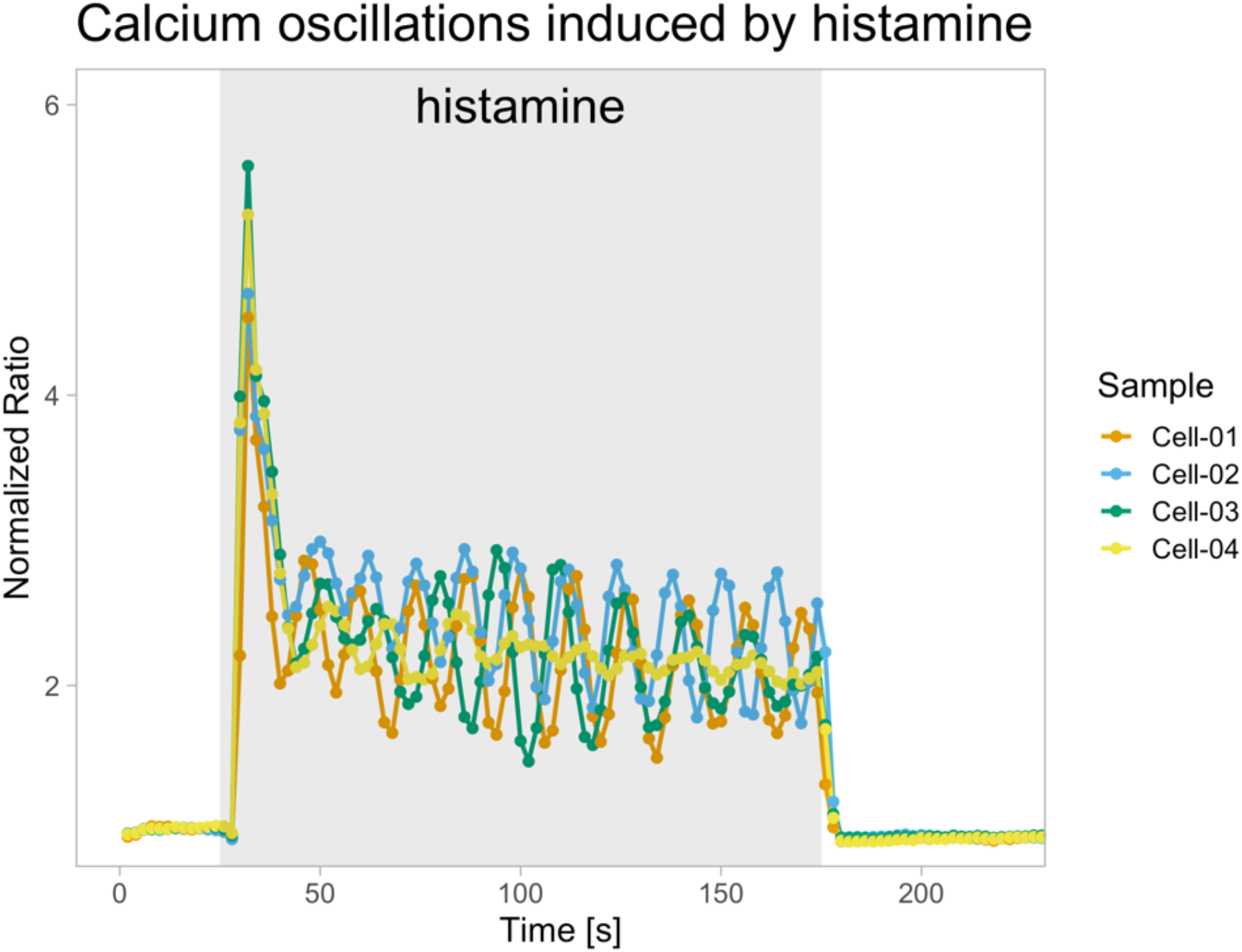
A plot that shows the change in YFP/CFP ratio induced by histamine over time. The ratio is used to indicate the intracellular calcium levels. The lines represent the data from four different cells, indicated with unique colors. The baseline fluorescence, determined from the average of the first 5 frames, was normalized 1.

- The script will produce a CSV file with normalized data that is saved in the working directory as ‘Normalized_data.csv’. This file can be used as input for PlotTwist: drop the file on the ‘Browse…’ button, select ‘These data are Tidy’ and as variable for the x-axis ‘Time’ and y-axis ‘ratio’ (or any of the other columns).
- Click on the ‘Plot’ tab to see the plot and adjust the visualization.
- To reproduce the plot, start PlotTwist with this URL: https://huygens.science.uva.nl:/PlotTwist/?data=3;TRUE;;fold;1,5;&vis=dataasline;1;;;1;TRUE;&layout=;;TRUE;0,220;;TRUE;;6;X;480;600&color=none&label=TRUE;Calciumoscillationsinducedbyhistamine;TRUE;Time[s];NormalizedRatio;TRUE;24;18;18;8;;;TRUE&stim=;bar;;;&
- Upload the file ‘Normalized_data.csv’, select variable for y-axis ‘ratio’ and click on the ‘Plot’ tab to show the plot.

**Figure 13:**
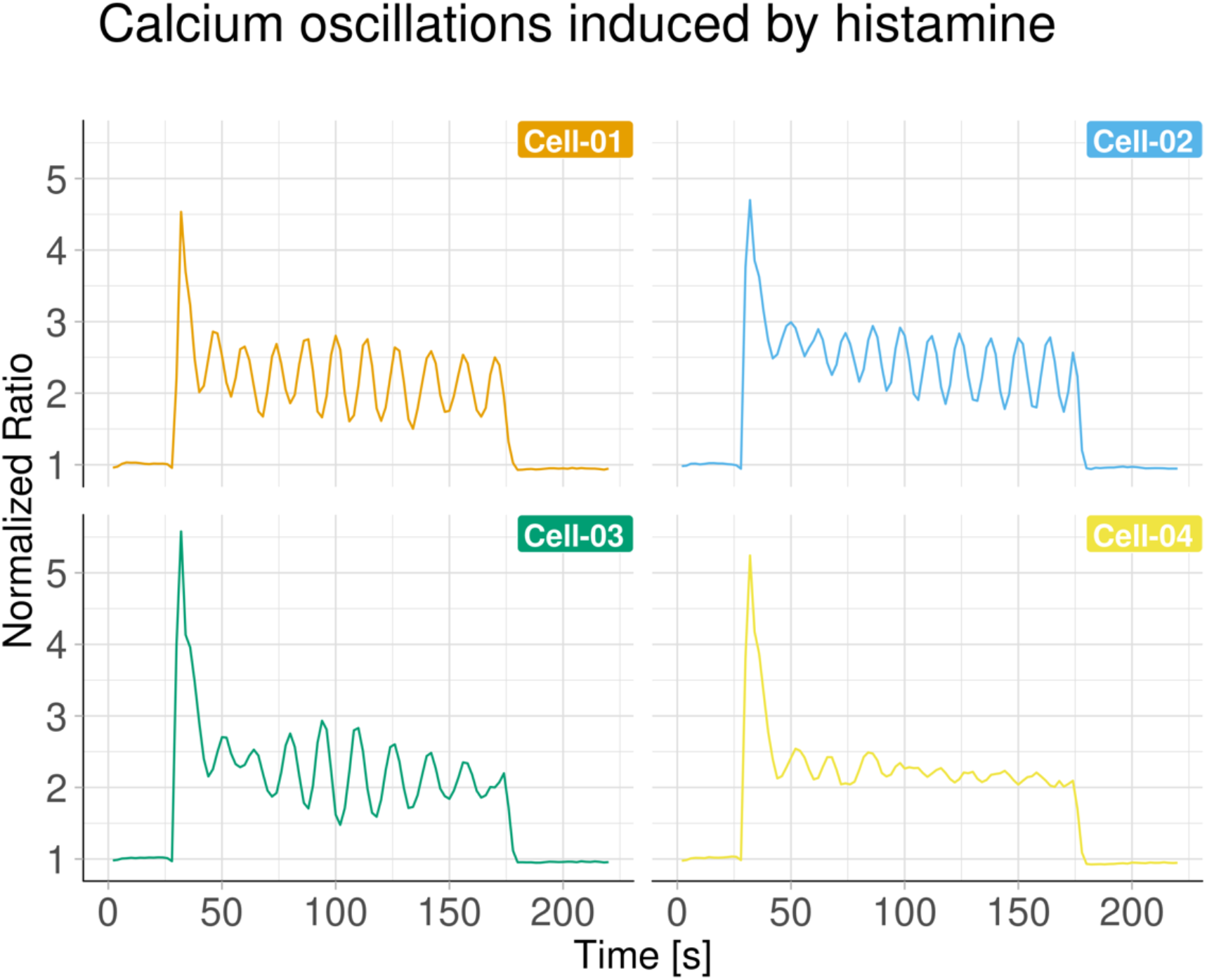
A plot that shows the change in YFP/CFP ratio induced by histamine over time. The ratio is used to indicate the intracellular calcium levels. The data from four different cells, indicated with unique colors, is shown as a small multiple. The baseline fluorescence, determined from the average of the first 5 frames, was normalized 1.

## 4 Notes

- Note 1: Use [Help > Search] to find the location of a function in a menu.
- Note 2: There are alternatives to open files: Alternative 1: [File > Open…] and locate the file on your computer. Alternative 2: [File > Import > URL…] and copy-past this URL in the text field: https://github.com/JoachimGoedhart/TimeLapse-Imaging-DataViz/raw/main/3_1_1_TemporalColor/201029_PMN_channel_06-2.tif Alternative 3: [File > Import > URL…] and copy-past this URL in the text field: https://zenodo.org/record/4501412/files/201029_PMN_channel_06-2.tif
- Note 3: When the function does not work, you may use the macro that is available on Github: https://github.com/JoachimGoedhart/TimeLapse-Imaging-DataViz/blob/main/Macros/Temporal-Color_Code.ijm See 2.2 for instructions on how to install the macro.
- Note 4: The choice of the LUT will influence the clarity of the visualization. The ‘Fire’ LUT that comes with FIJI is a safe first choice, but it is recommended to try a couple of different LUTs.
- Note 5: The position of the line can be stored by adding it in the ROI manager: [Analyze > Tools > ROI Manager…] and press the button ‘Add’. The line can be permanently saved by using [More > Save…] on the ROI manager.
- Note 6: KymoButler [15] is an online tool that quantifies track length, duration and velocity from a kymograph: https://www.wolframcloud.com/objects/deepmirror/Projects/KymoButler/KymoButlerForm The cloud app accepts a maximum of 150,000 pixels, so it makes sense to crop and analyze the upper half of the image.
- Note 7: This can also be done in other ways, for instance by [Edit > Copy] followed by [File > New > Internal Clipboard]
- Note 8: The results are in a ‘tidy format’, which implies that all the measured mean intensities are in a single column that is named ‘Mean’. Two other columns, ‘Label’ and ‘Slice’ hold information on the ROI and the slice number respectively. The tidy format is well-suited for handling the data with open-source application such as R and Python.
- Note 9: To run the entire script at once, select all the code and press <command>+<enter>.

## Acknowledgements

We thank Janine Arts and Jaap van Buul (Sanquin Research and Landsteiner Laboratory, Amsterdam, the Netherlands) for providing the data of the membrane dynamics and Marten Postma (University of Amsterdam, the Netherlands) for useful discussions. This work was supported by an NWO ALW-OPEN grant ALWOP.306 (EKM). We are grateful for all the input, comments and solutions from the active communities on stackoverflow, twitter and other fora that share their knowledge and expertise (you know who you are).

